# Contributions of local speech encoding and functional connectivity to audio-visual speech integration

**DOI:** 10.1101/097493

**Authors:** Bruno L. Giordano, Robin A. A. Ince, Joachim Gross, Stefano Panzeri, Philippe G. Schyns, Christoph Kayser

**Affiliations:** Institute of Neuroscience and Psychology, University of Glasgow, Glasgow, G12 8QB, UK; Institut de Neurosciences de la Timone UMR 7289, Aix-Marseille Université - Centre National de la Recherche Scientifique, 13005 Marseille, France; Neural Computation Laboratory, Center for Neuroscience and Cognitive Systems, Istituto Italiano di Tecnologia, Rovereto, 38068, Italy

## Abstract

Seeing a speaker’s face enhances speech intelligibility in adverse environments. We investigated the underlying network mechanisms by quantifying local speech representations and directed connectivity in MEG data obtained while human participants listened to speech of varying acoustic SNR and visual context. During high acoustic SNR speech encoding by entrained brain activity was strong in temporal and inferior frontal cortex, while during low SNR strong entrainment emerged in premotor and superior frontal cortex. These changes in local encoding were accompanied by changes in directed connectivity along the ventral stream and the auditory-premotor axis. Importantly, the behavioural benefit arising from seeing the speaker's face was not predicted by changes in local encoding but rather by enhanced functional connectivity between temporal and inferior frontal cortex. Our results demonstrate a role of auditory-motor interactions in visual speech representations and suggest that functional connectivity along the ventral pathway facilitates speech comprehension in multisensory environments.

## Introduction

When communicating in challenging acoustic environments we profit tremendously from visual cues arising from the speakers face. Movements of the lips, tongue or the eyes convey significant information that can boost speech intelligibility and facilitate the attentive tracking of individual speakers (Ross et al., 2007; Sumby and Pollack, 1954). This multisensory benefit is strongest for continuous speech, where visual signals provide temporal markers to segment words or syllables, and provide linguistic cues (Grant and Seitz, 1998). Previous work has identified the synchronization of brain rhythms between interlocutors as a potential neural mechanism underlying the visual enhancement of intelligibility (Hasson et al., 2012; Park et al., 2016; Peelle and Sommers, 2015; Pickering and Garrod, 2013; Schroeder et al., 2008). Both acoustic and visual speech signals exhibit pseudo-rhythmic temporal structures at prosodic and syllabic rates (Chandrasekaran et al., 2009). These regular features can entrain rhythmic activity in the observer’s brain and facilitate perception by aligning neural excitability with acoustic or visual speech features (Giraud and Poeppel, 2012; Mesgarani and Chang, 2012; Park et al., 2016; Peelle and Davis, 2012; Schroeder et al., 2008; van Wassenhove, 2013). While this model makes clear predictions about the visual enhancement of speech encoding in challenging environments, the network organization of multisensory speech enhancement remains unclear.

**Figure 1.**
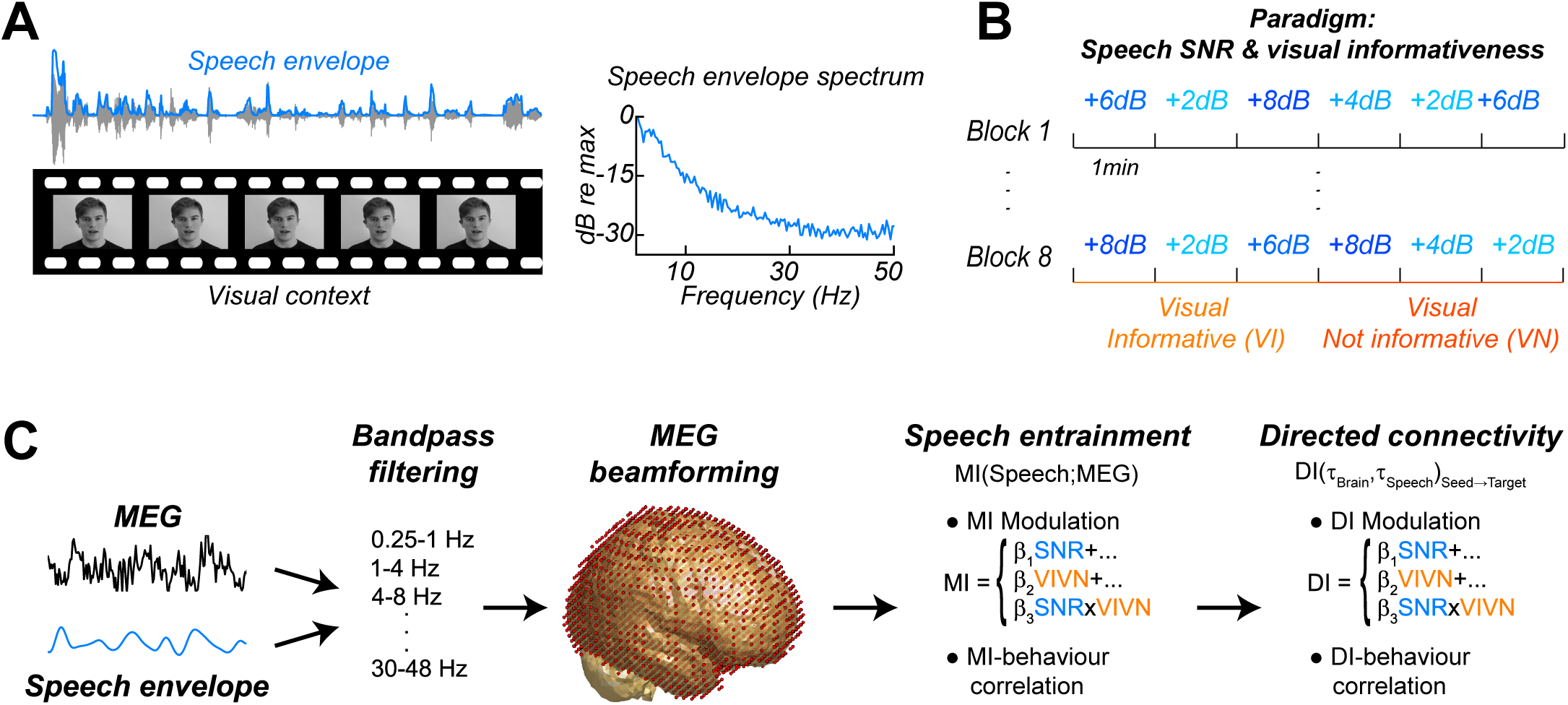
Experimental paradigm and analysis. **(A)** Stimuli consisted of 8 continuous 6 minute long audio-visual speech samples. **(B)** The experimental design comprised 8 conditions, defined by the factorial combination of 4 levels of speech to background signal to noise ratio (SNR = 2, 4, 6, and 8 dB) and two levels of visual informativeness (VI: Visual context Informative: video showing the narrator in synch with speech; VN: Visual context Not informative: video showing the narrator producing babble speech). Experimental conditions lasted 1 (SNR) or 3 (VIVN) minutes, and were presented in pseudo-randomized order. **(C)** Analyses were carried out on band-pass filtered speech envelope and MEG signals. The MEG data were source-projected onto a grey-matter grid (LCMV beamformer). One analysis quantified speech entrainment, i.e. the mutual information (MI) between the MEG data and the speech envelope, and the extent to which this was modulated by the experimental conditions. A second analysis quantified directed functional connectivity (DI) between seeds and the extent to which this was modulated by the experimental conditions. A final analysis assessed the correlation of either MI or DI with word-recognition performance.

Previous work has implicated many brain regions in the visual enhancement of speech, including superior temporal (Beauchamp et al., 2004; Nath and Beauchamp, 2011; Riedel et al., 2015; van Atteveldt et al., 2004), premotor and inferior frontal cortices (Arnal et al., 2009; Evans and Davis, 2015; Hasson et al., 2007b; Lee and Noppeney, 2011; Meister et al., 2007; Skipper et al., 2009; Wright et al., 2003). Furthermore, some studies have shown that the visual facilitation of speech encoding may even commence in early auditory cortices (Besle et al., 2008; Chandrasekaran et al., 2013; Ghazanfar et al., 2005; Kayser et al., 2010; Lakatos et al., 2009; Zion Golumbic et al., 2013). However, it remains to be understood whether visual context shapes the encoding of speech differentially within distinct regions of the auditory pathways, or whether the visual facilitation observed within auditory regions is simply fed forward to upstream areas, perhaps without further modification. Hence, it is still unclear whether the enhancement of speech-to-brain entrainment is a general mechanism that mediates visual benefits at multiple stages along the auditory pathways.

Many previous studies on this question were limited by three conceptual shortcomings: first, many have focused on generic brain activations rather than directly mapping the task-relevant sensory representations (activation mapping vs. information mapping, Kriegeskorte et al., 2006), and hence have not quantified multisensory influences on those neural representations directly shaping behavioural performance. Second, while many studies have correlated speech-induced local brain activity with behavioural performance, few studies have quantified directed connectivity along the auditory pathways to ask whether perceptual benefits are better explained by changes in local encoding or by changes in functional connectivity. And third, most studies have neglected the continuous predictive structure of speech by focusing on isolated words or syllables. However, this structure may play a central role for mediating the visual benefits (Bernstein et al., 2004; Giraud and Poeppel, 2012; Schroeder et al., 2008; Schwartz and Savariaux, 2014). Importantly, given that the predictive visual context interacts with acoustic signal quality to increase perceptual benefits in adverse environments (Callan et al., 2014; Ross et al., 2007; Schwartz et al., 2004; Sumby and Pollack, 1954), one needs to manipulate both factors to fully address this question. Overcoming these problems, we capitalized on the statistical and conceptual power offered by naturalistic speech to study the network mechanisms that underlie the visual facilitation of speech perception.

Using source localized MEG activity we systematically investigated how local speech representations and task-relevant directed functional connectivity along the auditory pathways change with visual context and acoustic signal quality. Specifically, we extracted neural signatures of speech representations by quantifying the mutual information between the MEG signal and the speech envelope. Furthermore, we quantified directed causal connectivity between nodes in the speech network using lagged mutual information between MEG source signals. Using linear modelling we then asked how local encoding and connectivity are affected by contextual information about the speakers face, by the acoustic signal to noise ratio, and by their interaction, and how each of these neural signatures relates to behavioural performance.

## Results

Participants (n = 19) were presented with continuous speech that varied in acoustic quality (signal to noise ratio, SNR) and the informativeness of the speaker’s face. The visual context could be either informative (VI), showing the face producing the acoustic speech, or uninformative (VN), showing the same face producing nonsense babble (Fig. 1A,B). We measured brain-wide activity using MEG while participants listened to eight six-minute texts and performed a delayed word recognition task. Behavioural performance was better during high SNR and an informative visual context (Fig. 2): a repeated measures ANOVA revealed a significant effect of SNR (F(3,54) = 36.22, p < 0.001, Huynh-Feldt corrected, 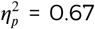), and of visual context (F(1,18) = 18.95, p < 0.001, 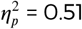),as well as a significant interaction (F(3,54) = 4.34, p = 0.008, 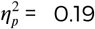). This interaction arose froma significant visual enhancement for SNRs of 4 and 8 dB (paired T(18) ≥ 3.00, Bonferroni corrected p ≤ 0.032; p > 0.95 for other SNRs).

**Figure 2.**
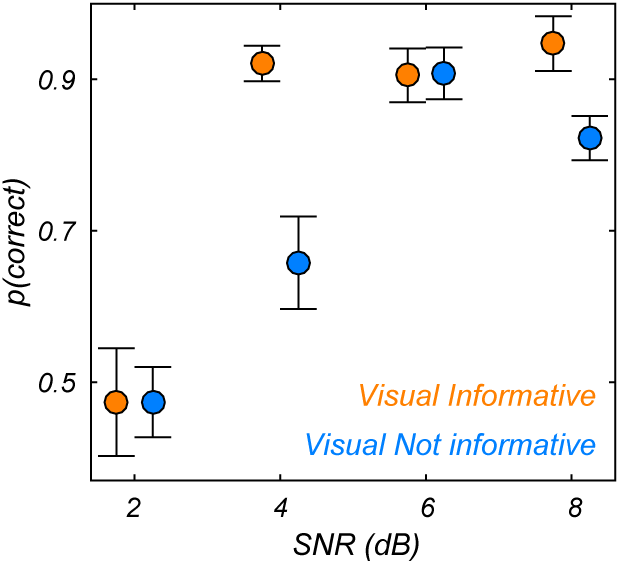
Behavioural performance. Word recognition performance for each of the experimental conditions (mean ± SEM across participants n=19).

To study the brain activity underlying this behavioral benefit we analyzed source-projected MEG data using information theoretic tools to quantify the fidelity of local neural representations of the speech envelope (speech-to-brain entrainment), as well as the directed causal connectivity between relevant regions. For both, coding and connectivity, we (1) modelled the extent to which they were modulated by the experimental conditions and (2) asked whether they correlated with behavioural performance across conditions and with the visual benefit (VI-VN) across SNRs (Fig. 1C).

### Widespread speech-to-brain entrainment at multiple time scales

Speech-to-brain entrainment was quantified by the mutual information (MI) between the MEG time course and the speech envelope (not the speech + noise mixture) in individual frequency bands (Gross et al., 2013; Kayser et al., 2015b, Fig. 1). At the group-level, we observed widespread significant speech MI in all considered bands from 0.25 to 48 Hz (FWE = 0.05), except between 18–24 Hz (Fig. S1A). Consistent with previous results (Gross et al., 2013; Ng et al., 2013; Park et al., 2016) speech MI was higher at low frequencies and strongest below 4 Hz (Fig. S1B). This time scale is typically associated with syllabic boundaries or prosodic stress (Giraud and Poeppel, 2012; Greenberg et al., 2003). Indeed, the average syllabic rate was 212 syllables per minute in the present material, corresponding to about 3.5 Hz. Across frequencies, MI was strongest in bilateral auditory cortex and more extended within the right hemisphere (Fig. S1B). Peak MI values were significantly higher in the right compared to the left hemisphere at frequencies below 12 Hz (paired t-tests; T(18) ≥ 3.1, p ≤ 0.043 Bonferroni corrected), and did not differ at higher frequencies (T(18) ≤ 2.78, p ≥ 0.08). Importantly, we observed significant speech-to-brain entrainment not only within temporal cortices but across multiple regions in the occipital, frontal and parietal lobes, consistent with the notion that speech information is represented also within motor and frontal regions (Bornkessel-Schlesewsky et al., 2015; Du et al., 2014; Skipper et al., 2009).

**Figure 3.**
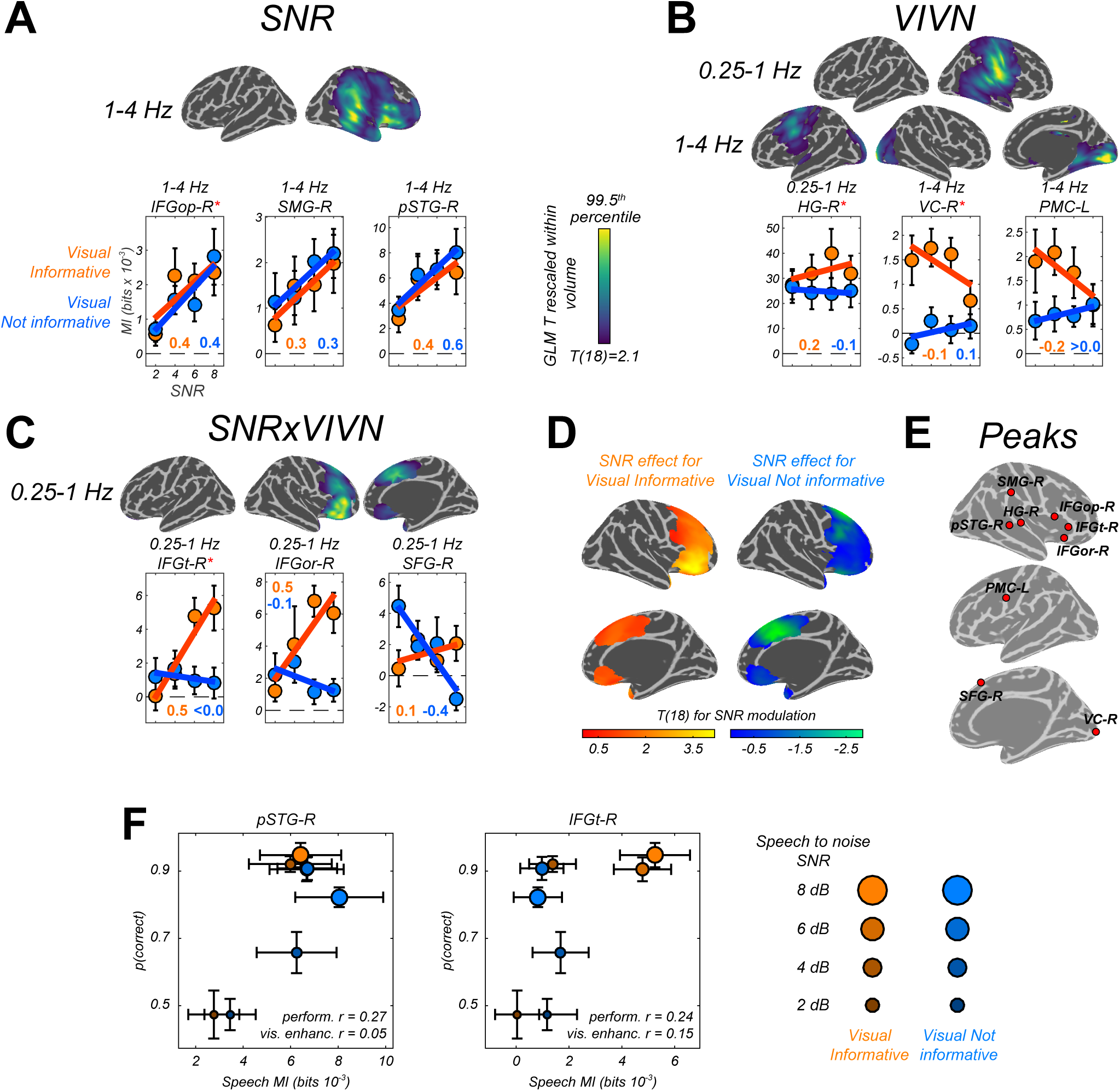
Modulation of speech-to-brain entrainment by acoustic SNR and visual informativeness. Changes in speech entrainment with the experimental factors were quantified using a GLM for the condition-specific speech MI based on the effects of SNR **(A)**, visual informativeness VIVN **(B)**, and their interaction (SNRxVIVN) **(C)**. The figures display the cortical-surface projection onto the Freesurfer template (proximity = 10 mm) of the group-level significant statistics for each GLM effect (FWE = 0.05). Graphs show the average speech MI values for each condition (mean ± SEM), for local and global (red asterisk) of the T maps. Lines indicate the across-participant average regression model and numbers indicate the group-average standardized regression coefficient for SNR in the VI and VN conditions. **(D)** T maps illustrating the opposite SNR effects within voxels with significant SNRxVIVN effects. MI graphs for the peaks of these maps are shown in (C) (IFGor-R and SFG-R = global T peaks for SNR effects in VI and VN, respectively). **(E)** Location of global and local seeds of GLM T maps, used for the analysis of directed connectivity. **(F)** Correlation between condition-specific behavioural performance and speech MI (perform. r) and between visual enhancement of performance and MI (vis. enhanc. r; see inset) in pSTG-R and IFGt-R. error-bars = ± SEM. See also Tables 1 and 3.

**Table 1.**
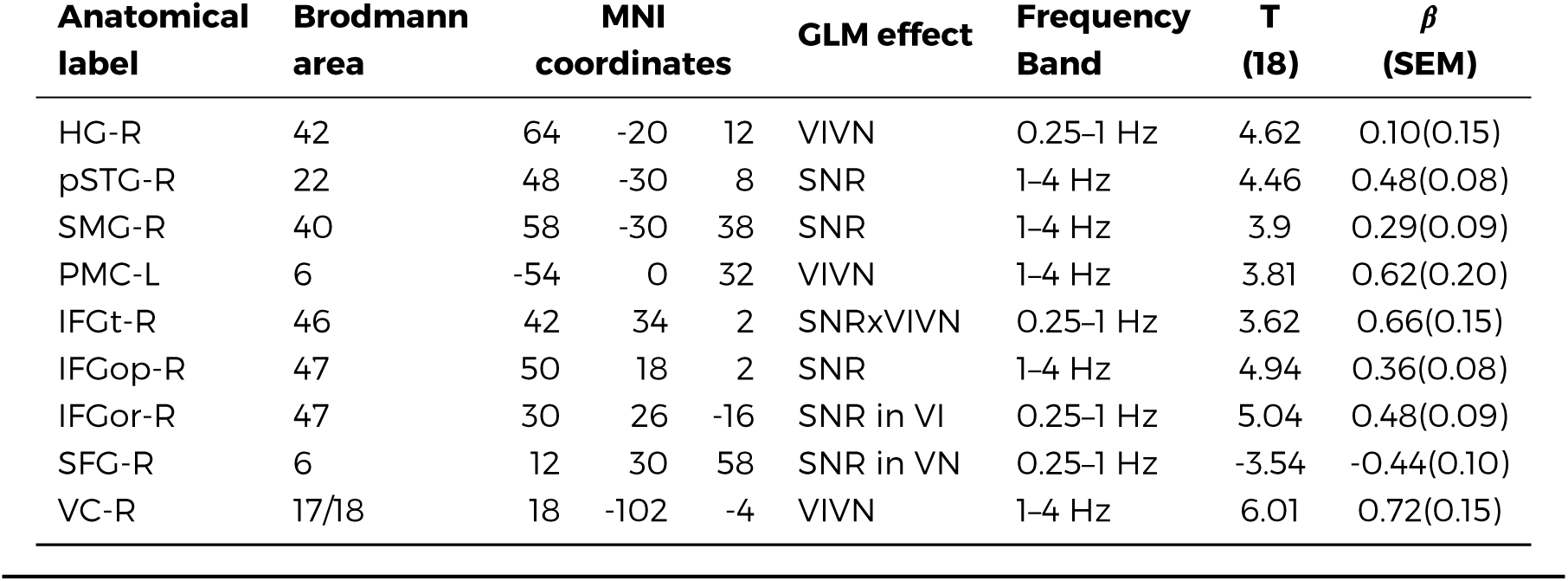
Condition effects on speech MI. The table lists global and local peaks in the GLM T-maps. Anatomical labels and Brodmann areas are based on the AAL and Talairach atlases. *β* = standardized regression coefficient; SEM = standard error of the participant average.

| Anatomical label | Brodmann area | MNI coordinates | | | GLM effect | Frequency Band | T (18) | $\beta$ (SEM) |
| --- | --- | --- | --- | --- | --- | --- | --- | --- |
| HG-R | 42 | 64 | -20 | 12 | VIVN | 0.25-1 Hz | 4.62 | 0.10(0.15) |
| pSTG-R | 22 | 48 | -30 | 8 | SNR | 1-4 Hz | 4.46 | 0.48(0.08) |
| SMG-R | 40 | 58 | -30 | 38 | SNR | 1-4 Hz | 3.9 | 0.29(0.09) |
| PMC-L | 6 | -54 | 0 | 32 | VIVN | 1-4 Hz | 3.81 | 0.62(0.20) |
| IFGt-R | 46 | 42 | 34 | 2 | SNRxVIVN | 0.25-1 Hz | 3.62 | 0.66(0.15) |
| IFGop-R | 47 | 50 | 18 | 2 | SNR | 1-4 Hz | 4.94 | 0.36(0.08) |
| IFGor-R | 47 | 30 | 26 | -16 | SNR in VI | 0.25-1 Hz | 5.04 | 0.48(0.09) |
| SFG-R | 6 | 12 | 30 | 58 | SNR in VN | 0.25-1 Hz | -3.54 | -0.44(0.10) |
| VC-R | 17/18 | 18 | -102 | -4 | VIVN | 1-4 Hz | 6.01 | 0.72(0.15) |

### Speech entrainment is modulated by SNR within and beyond auditory cortex

To determine the regions where acoustic signal quality and visual context affect the encoding of speech we modelled the condition-specific speech MI values based on effects of acoustic signal quality (SNR), visual informativeness (VIVN), and their interaction (SNRxVIVN). Random-effects significance was tested using a permutation procedure and cluster enhancement, correcting for multiple comparisons along all relevant dimensions. Effects of experimental factors emerged in multiple regions at frequencies below 4 Hz (Fig. 3). Increasing the acoustic signal quality (SNR; Fig. 3A) resulted in stronger speech MI in the right auditory cortex (1–4 Hz; local peakT statistic = 4.46 in posterior superior temporal gyrus; pSTG-R; Table 1), right parietal cortex (local peak T = 3.90 in supramarginal gyrus; SMG-R), and right dorsoventral frontal cortex (IFGop-R; global peak T = 4.94). We also observed significant positive SNR effects within the right temporo-parietal and occipital cortex at 12–18 Hz (local peak right lingual gyrus, T = 5.12). However, inspection of the participant-specific data suggested that this effect was not reliable (for only 58% of participants showed an speech MI increase with SNR, as opposed to a minimum of 84% for the other SNR effects), possibly because the comparatively lower power of speech envelope fluctuations at higher frequencies (c.f. Fig. 1A), and hence this effect is not discussed further.

### Visual context reveals distinct strategies for handling speech in noise in premotor, superior and inferior frontal cortex

Contrasting informative and un-informative visual contexts revealed stronger speech MI when seeing the speakers face (VI) at frequencies below 4 Hz in both hemispheres (Fig. 3B): the right temporoparietal cortex (0.25-1 Hz; HG;T = 4.62), bilateral occipital cortex (1–4 Hz; right visual cortex; VC-R; global T peak = 6.01) and left premotor cortex (1-4 Hz; PMC-L; local T peak = 3.81). Interestingly, the condition-specific pattern of MI for both VC-L and PMC-L were characterized by an increase in speech MI with decreasing SNR during the VI condition, pointing to a stronger visual enhancement during more adverse listening conditions.

Since visual benefits for perception emerge mostly when acoustic signals are degraded (Fig. 2, Ross et al., 2007; Sumby and Pollack, 1954), the interaction of acoustic and visual factors provides a crucial test for audio-visual integration. We found significant interactions in the 0.25–1 Hz band in the right dorsoventral frontal lobe, which peaked in the pars triangularis (IFGt-R; T = 3.62; Fig. 3C). Importantly, investigating the SNR effect at these voxels revealed two distinct strategies for handling speech in noise dependent on visual context (Fig. 3D): During VI speech MI increased with SNR in ventral frontal cortex (peakT for SNR in pars orbitalis; IFGor-R; T = 5.04), while in dorsal frontal cortex speech MI was strongest at low SNRs during VN (peakT in superior frontal gyrus; SFG-R; T = –3.54). This demonstrates distinct functional roles of ventral and dorsal prefrontal regions in speech encoding and reveals a unique role of superior frontal cortex for enhancing speech representations in a poorly informative context, such as the absence of visual information in conjunction with poor acoustic signals.

**Figure 4.**
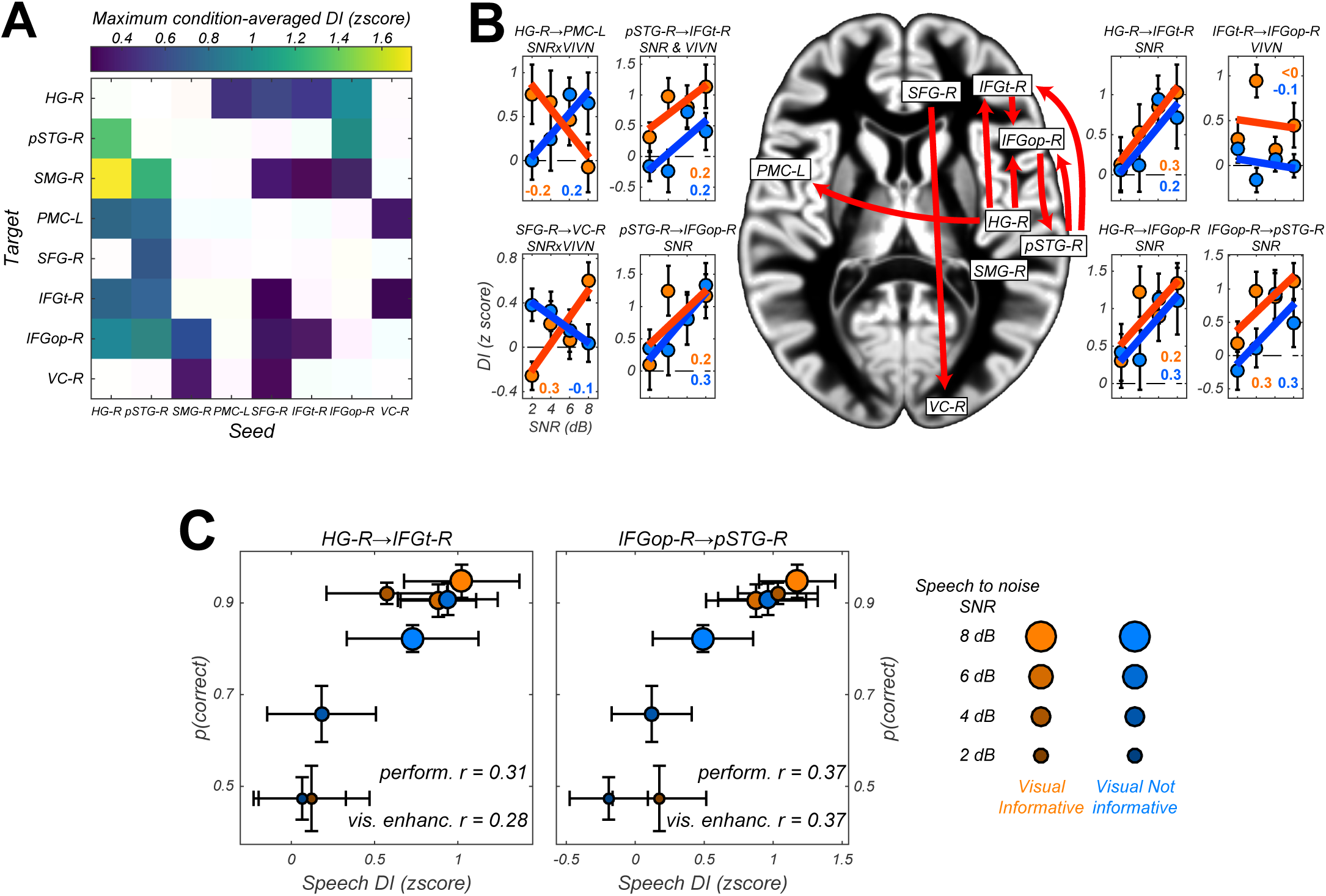
Directed causal connectivity within the speech-entrained network. Directed connectivity between seeds of interest (c.f. Fig. 3E) was quantified using Directed Information (DI). **(A)** Maximum significant condition-average DI across lags (FWE = 0.05 across lags; white = no significant DI). **(B)** Significant condition effects (GLM for SNR, VIVN or their interaction) on DI (FWE = 0.05 across speech/brain lags and seed/target pairs). Bar graphs display condition-specific DI values for each significant GLM effect along with the across-participants average regression model (lines). Numbers indicate the group-average standardized betas for SNR in the VI and VN conditions, averaged across lags associated with a significant GLM effect. **(C)** Correlation between behavioural performance and condition-specific DI (perform. r) and between visual enhancement of performance and DI (vis. enhanc. r) from HG-R to IFGt-R and from IFGop-R to pSTG-R. error-bars = ± SEM. See also Tables 2-3 and Fig. S2.

### Directed causal connectivity within the speech network

The diversity of the patterns of speech entrainment in temporal, premotor and inferior frontal regions across conditions could arise from the individual encoding properties of each region, or from changes in functional connectivity between regions with conditions. To directly test this, we quantified the directed causal connectivity between regions of interest extracted from the above analysis (Fig. 3E). To this end we used Directed Information (DI), also known as Transfer Entropy, an information theoretic measure of Wiener-Granger causality (Massey, 1990; Schreiber, 2000). We took advantage of previous work that made this measure statistically robust when applied to neural data (Besserve et al., 2015; Ince et al., 2016a).

**Table 2.** Analysis of directed connectivity (DI). The table lists connections with significant condition-averaged DI, and condition effects on DI. SEM = standard error of participant average; *β* = standardized regression coefficients. T(18) = maximum T statistic within significance mask.

| Seed | Target | DI<br>T(18) | Condition effects (GLM) |  |  |
| --- | --- | --- | --- | --- | --- |
| | | | Effect | T(18) | $\beta$(SEM) |
| HG-R | PMC-L | 3.38 | SNRxVIVN | -3.01 | -0.14(0.05) |
| HG-R | IFGt-R | 3.03 | SNR | 3.32 | 0.18(0.05) |
| HG-R | IFGopR | 4.54 | SNR | 3.19 | 0.18(0.05) |
| pSTG-R | IFGt-R | 3.39 | SNR | 3.91 | 0.22(0.06) |
|  |  |  | VIVN | 4.57 | 0.59(0.22) |
| pSTG-R | IFGopR | 4.12 | SNR | 3.31 | 0.20(0.06) |
| SFG-R | VC-R | 4.4 | SNRxVIVN | 3.69 | 0.12(0.03) |
| IFGt-R | IFGopR | 3.76 | VIVN | 3.56 | 0.31(0.17) |
| IFGopR | pSTG-R | 4.16 | SNR | 4.65 | 0.17(0.04) |

We observed significant condition-averaged DI between multiple nodes of the speech network (FWE = 0.05; Fig. 4A and Fig. S2A). This included among others the feed-forward pathways of the ventral and dorsal auditory streams, such as from auditory cortex (HG-R) and superior temporal regions (pSTG-R) to premotor (PMC-L) and to inferior frontal regions (IFGt-R, IFGop-R), from right parietal cortex (SMG-R) to premotor cortex (PMC-L), as well as feed-back connections from premotor and inferior frontal regions to temporal regions. In addition, we also observed significant connectivity between frontal (SFG-R) and visual cortex (VC).

We then asked whether and where connectivity changed with experimental conditions (Fig. 4B, Table 2 and Fig. S2B). Within the right ventral stream feed-forward connectivity from the temporal lobe (HG-R, pSTG-R) to frontal cortex (IFGt-R, IFGop-R) was enhanced during high acoustic SNR (FWE = 0.05; T(18) ≥ 3.1). More interestingly, this connectivity was further enhanced in the presence of an informative visual context (pSTG-R → IFGt-R, positive SNRxVIVN interaction, T = 4.57), demonstrating a direct influence of visual context on the propagation of speech information along the ventral stream. Interactions of acoustic and visual context on connectivity were also found from auditory (HG-R) to premotor cortex (PMC-L, negative interaction; T = –3.01). Here connectivity increased with increasing SNR in the absence of visual information and increased with decreasing SNR during an informative context, suggesting that visual information changes the qualitative nature of auditory-motor interactions. An opposite interaction was observed between the frontal lobe and visual cortex (SFG-R → VC-R, T = 4.40). Finally, we found that feed-back connectivity along the ventral pathway was significantly stronger during high SNRs (IFGt-R → pSTG-R; T = 4.16).

### Do Speech entrainment or connectivity shape behavioural performance?

We performed two additional analyses to test whether and where changes in the local representation of speech information (speech-MI) or directed connectivity (DI) contribute to explaining the behavioural benefits (Fig. 2). First, we asked where speech-MI/DI relates to performance changes across all experimental conditions (incl. changes in SNR). This revealed a significant correlation between condition-specific word-recognition performance and the strength of speech MI in pSTG-R and IFGt - R (r ≥ 0.28; FWE = 0.05; Table 3 and Fig. 3F), suggesting that stronger entrainment in the ventral stream facilitates comprehension. This hypothesis was further corroborated by a significant correlation of connectivity along the ventral stream with behavioural performance, both in feed-forward (HG-R → IFGt-R; pSTG-R → IFGt-R/IFGop-R; r > 0.27, Table 3) and feed-back directions (IFGop-R → pSTG-R; r=0.37). The enhanced quality of speech perception during favourable listening conditions hence results from enhanced speech encoding and the supporting network connections along the temporal-frontal axis.

**Table 3.** Association of behavioural performance with speech entrainment and connectivity. Performance: T statistic and average of participant-specific correlation (SEM) between behavioural performance and speech MI / DI. Visual enhancement: correlation between SNR-specific behavioural benefit (VI-VN) and the respective difference in speech-MI or DI. * FWE = 0.05 corrected for multiple comparisons.

| Speech MI |  |  |  |  |  |
| --- | --- | --- | --- | --- | --- |
|  |  | Performance |  | Visual enhancement |  |
|  |  | T(18) | r(SEM) | T(18) | r(SEM) |
| HG-R |  | 1.27 | 0.12(0.08) | 0.21 | 0.04(0.12) |
| pSTG-R |  | 3.43 * | 0.27(0.07) | 0.53 | 0.05(0.10) |
| SMG-R |  | 2.35 | 0.19(0.08) | -0.39 | -0.02(0.10) |
| PMC-L |  | 0.47 | 0.04(0.07) | 0.13 | 0.03(0.12) |
| SFG-R |  | -0.47 | -0.03(0.07) | 1.61 | 0.17(0.11) |
| IFGt-R |  | 3.09 * | 0.24(0.08) | 1.25 | 0.15(0.12) |
| IFGopR |  | 2.38 | 0.20(0.08) | -0.25 | -0.01(0.12) |
| VC-R |  | 1.55 | 0.14(0.09) | -0.82 | -0.16(0.11) |

| Directed connectivity |  |  |  |  |  |
| --- | --- | --- | --- | --- | --- |
|  |  | Performance |  | Visual enhancement |  |
| Seed | Target | T(18) | r(SEM) | T(18) | r(SEM) |
| HG-R | IFGt-R | 4.83 * | 0.31(0.07) | 2.55 * | 0.28(0.11) |
| HG-R | IFGopR | 3.19 * | 0.24(0.07) | 1.86 | 0.31(0.17) |
| HG-R | PMC-L | 0.90 | 0.06(0.06) | -0.07 | -0.01(0.14) |
| pSTG-R | IFGt-R | 4.28 * | 0.27(0.06) | 1.28 | 0.16(0.12) |
| pSTG-R | IFGopR | 3.59 * | 0.29(0.08) | 1.82 | 0.32(0.17) |
| IFGt-R | IFGopR | 1.11 | 0.08(0.07) | 2.27 | 0.33(0.14) |
| IFGopR | pSTG-R | 4.51 * | 0.37(0.08) | 2.55 * | 0.37(0.15) |
| SFG-R | VC-R | -0.04 | 0.00(0.08) | 0.90 | 0.17(0.18) |

Second, we asked whether and where the improvement in behavioural performance with an informative visual context (VI-VN) correlates with an enhancement in speech encoding or connectivity. This revealed no significant correlation between the visual enhancement of local speech representations and perceptual benefits (all p > 0.05). However, both feed-forward (HG-R → IFGt-R; r= 0.28, p < 0.05; Fig. 4C) and feedback connections (IFGop-R → pSTG-R; r = 0.37) along the ventral stream were significantly enhanced during an informative visual context, suggesting that changes in functional connectivity contribute significantly to shaping speech intelligibility.

## Discussion

The present study provides a comprehensive picture of how acoustic signal quality and visual context interact to shape the encoding of speech information and the directed functional connectivity along speech-sensitive cortex. Our results reveal a dominance of feed-forward pathways from auditory regions to inferior frontal cortex under favourable conditions, such as during high SNR. We also demonstrate the visual enhancement of speech encoding in auditory and premotor cortex, as well as non-trivial interactions of acoustic quality and visual context in superior and inferior frontal regions. These patterns of local encoding were accompanied by changes in directed connectivity along the ventral pathway and from auditory to premotor cortex. Yet, the behavioural benefit arising from seeing the speaker’s face was not related to any site-specific visual enhancement of local speech encoding. Rather, changes in directed functional connectivity along the ventral stream were predictive of the behavioural benefit.

### Entrained speech representations in temporal, parietal and frontal lobes

We observed functionally distinct patterns of speech-to-brain entrainment along the auditory pathways. Previous studies on speech entrainment largely focused on the auditory cortex, where entrainment is strongest (Ding and Simon, 2013; Gross et al., 2013; Keitel et al., 2011; Mesgarani and Chang, 2012). This was in part due to the difficulty to separate distinct processes reflecting entrainment when contrasting only few experimental conditions (e.g. forward and reversed speech, Ding and Simon, 2012; Gross et al., 2013). Based on the susceptibility to changes in acoustic signal quality and visual context we here establish entrainment as a ubiquitous mechanism reflecting distinct speech representations along auditory pathways.

Speech entrainment was reduced with decreasing acoustic SNR in temporal, parietal and ventral prefrontal cortex, directly reflecting the reduction in behavioural performance in challenging environments. In contrast, entrainment was enhanced during low SNR in superior frontal and premotor cortex. While there is strong support for a role of frontal and premotor regions in speech analysis (Du et al., 2014; Evans and Davis, 2015; Heim et al., 2008; Meister et al., 2007; Morillon et al., 2015; Rauschecker and Scott, 2009; Skipper et al., 2009; Wild et al., 2012), most evidence comes from stimulus-evoked activity rather than signatures of neural speech encoding. We directly demonstrate the specific enhancement of frontal (PMC, SFG) speech representations during challenging conditions. This enhancement is not directly inherited from the temporal lobe, as temporal regions exhibited either no visual facilitation (pSTG) or visual facilitation without an interaction with SNR (HG).

The effects of experimental conditions dominated on the right hemisphere. Such a right dominance of speech entrainment is in agreement with previous studies (Bourguignon et al., 2013; Fonteneau et al., 2015; Gross et al., 2013; Vander Ghinst et al., 2016) and with the hypothesis that right temporal regions extract acoustic information predominantly on the syllabic and prosodic time scales (Giraud and Poeppel, 2012; Poeppel, 2003), exactly those timescaleswherespeech-to-brain entrainment is strongest in the present and previous data (Gross et al., 2013; Keitel et al., 2017).

### Multisensory enhancement of speech encoding in the frontal lobe

Visual information from the speakers face provides multiple cues that enhance intelligibility. In support of a behavioural multisensory benefit we found stronger entrainment during an informative visual context in multiple bilateral regions. First, we replicated the visual enhancement of auditory cortical representations (HG, Besle et al., 2008; Kayser et al., 2010; Zion Golumbic et al., 2013).Second, visual enhancement of an acoustic speech representation was also visible in early visual areas, as suggested by prior studies (Nath and Beauchamp, 2011; Schepers et al., 2015). While we can’t rule out that this effect is in part mediated by the correlations between acoustic and visual speech cues, we found that the visual enhancement was strongest when SNR was low and hence is better explained by top-down influences (Vetter et al., 2014). Third, speech representations in ventral prefrontal cortex were selectively involved during highly reliable multisensory conditions and were reduced in the absence of the speakers face. These findings are in line with suggestions that the IFG facilitates comprehension (Alho et al., 2014; Evans and Davis, 2015; Hasson et al., 2007b; Hickok and Poeppel, 2007) and implements multisensory processes (Callan et al., 2014, 2003; Lee and Noppeney, 2011), possibly by providing amodal phonological, syntactic and semantic processes (Clos et al., 2014; Ferstl et al., 2008; McGettigan et al., 2012). Previous studies often reported enhanced IFG response amplitudes under challenging conditions (Guediche et al., 2013). In contrast, by quantifying the fidelity of speech representations we here show that these are generally stronger during favourable SNRs. This discrepancy is not necessarily surprising, if one assumes that IFG representations are derived from those in the temporal lobe, which are also more reliable during high SNRs. Noteworthy, however, is the finding that representations within ventral IFG are selectively stronger during an informative visual context. We thereby directly confirm the hypothesis that IFG speech encoding is enhanced by visual context.

Furthermore, we demonstrate the visual enhancement of speech representations in premotor regions, which could implement the mapping of audio-visual speech features onto articulatory representations (Meister et al., 2007; Morillon et al., 2015; Fernández et al., 2015; Skipper et al., 2009; Wilson et al., 2004). We show that that this enhancement is inversely related to acoustic signal quality. While this observation is in agreement with the notion that perceptual benefits are strongest under adverse conditions (Ross et al., 2007; Sumby and Pollack, 1954), there was no significant correlation between the visual enhancement of premotor encoding and behavioural performance. Our results thereby deviate from previous work that has suggested a driving role of premotor regions in shaping intelligibility (Alho et al., 2014; Osnes et al., 2011), and we rather support a modulatory influence of auditory-motor interactions (Alho et al., 2014; Callan et al., 2004; Hickok and Poeppel, 2007; Krieger-Redwood et al., 2013; Morillon et al., 2015). For example, in a study quantifying dynamic representations of visual speech signals (lip movements) we recently found that left premotor activity was significantly predictive of behavioural performance (Park et al., 2016). One explanation for this discrepancy may be presence of a memory component in our behavioural task, which may engage other brain regions (e.g. IFG) more than other tasks. Given that the premotor effects were restricted to the theta band, which is associated with syllabic (> 3 Hz) rather than intonational (< 1 Hz) structure (Giraud and Poeppel, 2012; Greenberg et al., 2003), our results also suggest this region carries syllabic rather than prosodic representations (Du et al., 2014; Heim et al., 2008; Krieger-Redwood et al., 2013; Osnes et al., 2011).

Finally, our results highlight an interesting role of the superior frontal gyrus, where entrainment was s’trongest when sensory information was most impoverished (low SNR, visual not informative) or when the speakers face was combined with clear speech (high SNR, visual informative). Superior frontal cortex has been implied in high level inference processes underlying comprehension, sentence level integration or the exchange with memory (Ferstl et al., 2008; Hasson et al., 2007a; Yarkoni et al., 2008) and is sometimes considered part of the broader semantic network (Binder et al., 2009; Gow and Olson, 2016; Price, 2012). Our data show that the SFG plays a critical role for speech encoding under challenging conditions at the supra-syllabic time scale, possibly by mediating sentence-level integration during low SNRs or the comparison of visual prosody with acoustic inputs in multisensory contexts.

### Multisensory behavioural benefits arise from distributed network mechanisms

To understand whether the condition-specific patterns of local speech representations emerge within each region, or whether they are possibly established by network interactions we investigated the directed functional connectivity between regions of interest. While many studies have assessed the connectivity between auditory regions (e.g. Abrams et al., 2013; Chu et al., 2013; Fonteneau et al., 2015; Park et al., 2015), few have quantified the behavioural relevance of these connections (Alho et al., 2014)

We observed significant intra-hemispheric connectivity between right temporal, parietal and frontal regions, in line with the transmission of speech information from auditory cortices along the auditory pathways (Bornkessel-Schlesewsky et al., 2015; Hickok, 2012; Poeppel, 2014). Supporting the idea that acoustic representations are progressively transformed along these pathways we found that the condition-specific patterns of functional connectivity differed systematically along the ventral and dorsal streams. While connectivity along the ventral stream was predictive of behavioural performance and strongest during favourable listening conditions, the inter-hemispheric connectivity to left premotor cortex was strongest during adverse multisensory conditions. Our results therefore suggest that premotor representations are informed by auditory regions (HG, pSTG) rather than being driven by the frontal lobe, an interpretation that is supported by previous work (Alho et al., 2014; Gow and Olson, 2016; Osnes et al., 2011).

Across changes in visual context and acoustic SNR behavioural performance was supported both by an enhancement of speech representations along multiple regions of the ventral pathway and increases in their functional connectivity. These increases in functional connectivity emerged both along feed-forward and feed-back directions between temporal and inferior frontal regions, and were strongest (in effect size) along the feed-back route. This underlines the hypothesis that recurrent processing, rather than a simple feed-forward sweep, is central to speech intelligibility (Bornkessel-Schlesewsky et al., 2015; Hickok, 2012; Poeppel, 2014). Central to the scope of the present study, however, we found that no single region-specific effect could explain the visual behavioural benefit. Rather, the benefit arising from seeing the speakers face was significantly correlated with the enhancement of functional connectivity along the ventral stream (HG → IFG → pSTG). Our results hence point to a distributed origin of the visual enhancement of speech intelligibility. As proposed a decade ago (Besle et al., 2008; Ghazanfar et al., 2005; Ghazanfar and Schroeder, 2006; Kayser et al., 2010; Zion Golumbic et al., 2013) this visual enhancement involves early auditory cortices, but as we show here, the behavioural benefit also relies on the recurrent transformation of speech representations between temporal and frontal regions. One interpretation of this in the context of predictive coding models is that an informative visual context facilitates the correction of prior predictions about the expected stimulus by incoming sensory evidence, which would be visible both in feed-forward and feed-back connectivity (Arnal and Giraud, 2012; Bastos et al., 2012).

Our results provide a network view on the dynamic speech representations in multisensory environments. While premotor and superior frontal regions are specifically engaged in the most challenging environments the visual enhancement of comprehension at intermediate SNRs is mediated by interactions of the core speech regions along the ventral pathway. Such a distributed neural origin of multisensory benefits is in line with the notion of a hierarchical organization of multisensory processing in the brain (Lee and Noppeney, 2011; Rohe and Noppeney, 2015), and the idea that comprehension is shaped by network connectivity more than the engagement of particular brain regions (Abrams et al., 2013).

## Materials and methods

Nineteen right handed healthy adults (10 females; age from 18 to 37) participated in this study. All participants were tested for normal hearing, were briefed about the nature and goal of this study, and received financial compensation for their participation. The study was conducted in accordance with the Declaration of Helsinki and was approved by the local ethics committee (College of Science and Engineering, University of Glasgow). Written informed consent was obtained from all participants.

### Stimulus material

The stimulus material consisted of audio-visual recordings based on text transcripts taken from publicly available TED talks also used in a previous study (Kayser et al., 2015b, Fig. 1A). Acoustic (44.1 kHz sampling rate) and video recordings (25 Hz frame rate, 1920 by 1080 pixels) were obtained while a trained male native English speaker narrated these texts (Kayser et al., 2015a). The root mean square (RMS) intensity of each audio recording was normalized using 6 s sliding windows to ensure a constant average intensity. Across the eight texts the average speech rate was 160 words (range 138–177) per minute, and the syllabic rate was 212 syllables (range 192–226) per minute.

### Experimental design and stimulus presentation

We presented each of the eight texts as continuous 6 minute sample, while manipulating the acoustic quality and the visual relevance in a block design within each text (Fig. 1B). The visual relevance was manipulated by either presenting the video matching the respective speech (visual informative, VI) or presenting a 3 s babble sequence that was repeated continuously (visual not informative, VN), and which started and ended with the mouth closed to avoid transients. The signal to noise ratio (SNR) of the acoustic speech was manipulated by presenting the speech on background cacophony of natural sounds and scaling the relative intensity of the speech while keeping the intensity of the background fixed. We used relative SNR values of+8, +6, +4 and +2 dB RMS intensity levels. The acoustic background consisted ofa cacophony of naturalistic sounds, created by randomly superimposing various naturalistic sounds from a larger database (using about 40 sounds at each moment in time, Kayser et al., 2016). This resulted in a total of 8 conditions (four SNR levels; visual informative or irrelevant) that were introduced in a block design (Fig. 1B). The SNR changed from minute to minute in a pseudo-random manner (12 one minute blocks per SNR level). Visual relevance was manipulated within 3 minute sub-blocks. Texts were presented with self-paced pauses. Subjects performed a delayed comprehension tasks after each block, whereby they had to indicate whether a specific word (noun) was mentioned in the previous text (6 words per text) or not (6 words per text) in a two alternative forced choice task. The words chosen from the presented text were randomly selected and covered all eight conditions. The average performance was 73±2% correct (mean and SEM across subjects), showing that they indeed paid attention to the stimulus. Behavioural performance was averaged within each condition, and analysed using a repeated measures ANOVA, with SNR and VIVN as within-subject factors. The stimulus presentation was controlled using the Psychophysics toolbox in Matlab (Brainard, 1997). Acoustic stimuli were presented using an Etymotic ER-30 tube-phone (tube length = 4 m) at 44.1 kHz sampling rate and an average intensity of 65 dB RMS level, calibrated separately for each ear. Visual stimuli were presented in grey-scale and projected onto a translucent screen at 1280 x 720 pixels at 25 fps covering a field of view of 41 x 33 degrees.

### Pre-processing of the speech envelope

We extracted the envelope of the speech signal (not the speech plus background mixture) by computing the wide-band envelope at 150 Hz temporal resolution as in previous work (Chandrasekaran et al., 2009; Kayser et al., 2015b). The speech signal was filtered (4^th^ order Butterworth filter; forward and reverse) into six frequency bands (100 Hz-4 kHz) spaced to cover equal widths on the cochlear map. The wide-band envelope was defined as the average of the Hilbert envelopes of these band-limited signals (c.f. Fig. 1A).

### MEG data collection

MEG recordings were acquired with a 248-magnetometers whole-head MEG system (MAGNES 3600 WH, 4-D Neuroimaging) at a sampling rate of 1017.25 Hz. Participants were seated upright. The position of five coils, marking fiducial landmarks on the head of the participants, was acquired at the beginning and at the end of each block. Across blocks, and participants, the maximum change in their position was 3.6 mm, on average (STD = 1.2 mm).

### MEG pre-processing

Analyses were carried out in Matlab using the Fieldtrip toolbox (Oostenveld et al., 2010), SPM12, and code for the computation of information-theoretic measures (Ince et al., 2016a). Block-specific data were pre-processed separately. Infrequent SQUID jumps (observed in 1.5% of the channels, on average) were repaired using piecewise cubic polynomial interpolation. Environmental magnetic noise was removed using regression based on principal components of reference channels. Both the MEG and reference data were filtered using a forward-reverse 70 Hz FIR low-pass (−40 dB at 72.5 Hz); a 0.2 Hz elliptic high-pass (−40 dB at 0.1 Hz); and a 50 Hz FIR notch filter (−40 dB at 50 ± 1Hz). Across participants and blocks, 7 MEG channels were discarded as they exhibited a frequency spectrum deviating consistently from the median spectrum (shared variance < 25%). For analysis signals were resampled to 150 Hz, high-pass filtered at 0.2 Hz (forward-reverse elliptic filter). ECG and EOG artefacts were removed using ICA in fieldtrip (runica on 40 principal components), and were identified based on the time course and topography of IC components (Hipp and Siegel, 2013).

### Structural data and source localization

High resolution anatomical MRI scans were acquired for each participant (voxel size = 1 mm^3^) and co-registered to the MEG data using a semi-automated procedure. Anatomicals were segmented into grey and white matter and cerebro-spinal fluid (Ashburner and Friston, 2005). The parameters for the affine registration of the anatomical to the MNI template were estimated, and used to normalize the grey matter probability maps of each individual to the MNI space. A group MNI source-projection grid with a resolution of 3 mm was prepared including only voxels associated with a group-average grey-matter probability of at least 0.25. The projection grid excluded various subcortical structures, identified using the AAL atlas (e.g., vermis, caudate, putamen and the cerebellum). Leadfields were computed based on a single shell conductor model. Time-domain projections were obtained on a block-by-block basis for each frequency band using LCMV spatial filters (regularization = 5%) along the dipole orientation of maximum variance.

### Analysis of speech to brain entrainment

Motivated by previous work (Gross et al., 2013; Ng et al., 2013), we considered eight partly overlapping frequency bands (0.25–1 Hz, 1–4 Hz, 4–8 Hz, 8–12 Hz, 12–18 Hz, 18–24 Hz, 24–36 Hz, and 30–48 Hz), and isolated them from the full-spectrum MEG and speech envelope signals using a forward-reverse 4^th^ order Butterworth filter (magnitude of frequency response at band limits = –6 dB). Entrainment was quantified using the mutual information (MI) between the filtered MEG and speech-envelope time courses (Cogan and Poeppel, 2011; Gross et al., 2013; Kayser et al., 2015b; Keitel et al., 2017; Ng et al., 2012). The MI was calculated using a recently developed binless approach based on statistical copulas, which provides greater sensitivity than methods based on binned signals (Ince et al., 2016a).

To quantify the entrainment of brain activity to the speech envelope we first determined the optimal time lag between MEG signals and the acoustic stimulus for individual bands and source voxels using a permutation-based RFX estimate. Lag estimates were obtained based on a quadratic fit, excluding lags with insignificant MI (permutation-based FDR = 0.01). Voxels without an estimate were assigned the median estimate within the same frequency band, and volumetric maps of the optimal lags were smoothed with a Gaussian (FWHM = 10 mm). Speech MI was then estimated for each band and voxel using the optimal lag. The significance of group-level speech MI assessed within a permutation-based RFX framework that relied on MI values corrected for bias at the single-subject level, and on cluster mass enhancement of the test statistics corrected for multiple comparisons at the second level (Maris and Oostenveld, 2007). At the single-subject level, null distributions were obtained by shuffling the assignment of speech and MEG, independently for each participant, i.e. by permuting the 6 speech segments within each of the 8 experimental conditions (using the same permutation across bands). Participant-specific bias-corrected speech MI values were then defined as the actual MI minus the median MI across all 720 possible null permutations. Group-level RFX testing relied on T-statistics for the null-hypothesis that the participant-averaged bias-corrected MI was significantly larger than zero. To this end we generated 10,000 samples of the group-averaged MI from the participant-specific null distributions, used cluster-mass enhancement across voxels and frequencies (cluster-forming threshold T(18) = 2.1) to extract the maximum cluster T across frequency bands and voxels, and considered as significant a cluster-enhanced T statistic higher than the 95^th^ percentile of the permutation distribution (corresponding to FWE = 0.05).

To determine whether speech entrainment was modulated by the experimental factors we used a permutation-based RFX GLM framework (Winkler et al., 2014). For each participant individually we considered the condition-specific bias-corrected MI averaged across repetitions and estimated the coefficients of a GLM for predicting MI based on SNR (2, 4, 6, 8 dB), VIVN (1 = Visual Informative; −1 = Visual Not informative), and their interaction. We computed a group-level T-statistic for assessing the hypothesis that the across-participant average GLM coefficient was significantly different than zero, using cluster-mass enhancement across voxels and frequencies. Permutation testing relied on the Freedman-Lane procedure (Freedman and Lane, 1983). Independently for each participant and GLM effect, we estimated the parameters of a reduced GLM that includes all of the effects but the one to be tested and extracted the residuals of the prediction. We then permuted the condition-specific residuals and extracted the GLM coefficient for the effect of interest estimated for these reshuffled residuals. We obtained a permutation T statistic for the group-average GLM coefficient of interest using the max-statistics. We considered as significant T values whose absolute value was higher than the 95^th^ percentile of the absolute value of 10,000 permutation samples, correcting for multiple comparisons across voxels / bands (FWE = 0.05). We only considered significant GLM effects in conjunction with a significant condition-average entrainment.

### Analysis of directed functional connectivity

To quantify directed functional connectivity we relied on the concept of Wiener-Granger causality and its information theoretic implementation known as Transfer Entropy or directed information (DI, Massey, 1990; Schreiber, 2000; Vicente et al., 2011; Wibral et al., 2011). Directed information in its original formulation (Massey, 1990, termed DI* here) quantifies causal connectivity by measuring the degree to which the past of a seed predicts the future of a target signal, conditional on the past of the target, defined at a specific lag (*τ_Brain_*):

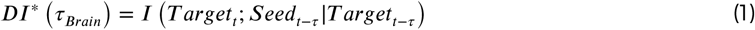

While DI* provides a measure of the overall directed influence from seed to target, it can be susceptible to statistical biases arising from limited sampling, common inputs or signal auto-correlations (Besserve et al., 2015, 2010; Ince et al., 2016a; Panzeri et al., 2007). We regularized and made this measure more conservative by subtracting out values of DI computed at fixed values of speech envelope. This subtraction removes terms – including the statistical biases described above – that cannot possibly carry speech information (because they are computed at fixed speech envelope). This results in an estimate that is statistically more robust, more conservative and more directly related to changes in the sensory input than classical transfer entropy (termed directed feature information in Ince et al., 2015, 2016a). Practically, DI was defined here as

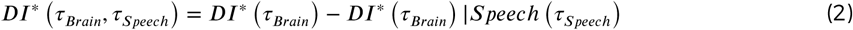

where DI\****|Speech*** denotes the DI* conditioned on the speech envelope. Positive values of DI indicate directed functional connectivity between seed and target at a specific brain (*τ_Brain_*) and speech lag (*τ_Speech_*) The actual DI values were furthermore Z-scored against random effects to further enhance the robustness of this connectivity index, which facilitates statistical comparisons between conditions across subjects (Besserve et al., 2015). To this end DI, as estimated for each participant and connection from Eq. 2, was Z-scored against the distribution of DI values obtained from condition-shuffled estimates (using the same randomization procedure as for MI). DI was computed for speech lags between 0 and 500 ms and brain lags between 0 and 250 ms, at steps of one sample (1/150 Hz). We estimated DI on the frequency range of 0.25-8 Hz (forward-reverse 4th order Butterworth filter) and by considering the bivariate MEG response defined by the band-passed source signal and its first-order difference (Ince et al., 2016a,b). Seeds for the DI analysis were the global and local peaks of the GLM-T maps quantifying the SNR, VIVN and SNRxVIVN modulation of entrainment, and the SFG-R voxel characterized by the peak negative effect of SNR in the visual informative condition, for a total of 8 seeds (Table 1 and Fig. 3E). To test for the significance of condition-average DI we used the same permutation-based RFX approach as for speech MI, testing the hypothesis that bias-corrected DI > 0. We used 2D cluster-mass enhancement of the T statistics within speech/brain lag dimensions correcting for multiple comparisons across speech and brain lags (FWE = 0.05). To test for significant DI effects with experimental conditions we relied on the same GLM strategy as for MI effects, again with the same differences pertaining to cluster enhancement and comparison correction (FWE = 0.05 across lags and seed/target pairs). We only considered DI modulations in conjunction with a significant condition-average DI.

### Neuro-behavioural correlations

We used a permutation-based RFX approach to assess (1) whether an increase in condition-specific speech-MI or DI was associated with an increase in behavioural performance, and (2) whether the visual enhancement (VI-VN) of MI or DI was associated with stronger behavioural gains. We focused on the 8 regions used as seeds for the DI analysis. For speech-MI we initially tested whether the participant-average Fisher Z-transformed correlation between condition-specific performance and speech-MI was significantly larger than zero. Uncorrected p-values were computed using the percentile method, where FWE = 0.05 p-values corrected across regions were computed using maximum statistics. We subsequently tested the positive correlation between SNR-specific visual gains (VI-VN) in speech-MI and behavioural performance using the same approach, but considered only those regions characterized by a significant condition-specific MI/performance association. For DI, we focused on those lags characterized by a significant SNR, VIVN, or SNRxVIVN DI modulation. Significance testing proceeded as for speech MI, except that Z-transformed correlations were computed independently for each lag and then averaged across lags (FWE = 0.05 corrected across all seed/target pairs).

## Acknowledgements

This research was supported by the UK Biotechnology and Biological Sciences Research Council (BBSRC, BB/L027534/1). CK is supported by the European Research Council (ERC-2014-CoG; grant No 646657); BLG by BBSRC BB/M009742/1; JG by the Wellcome Trust (Joint Senior Investigator Grant, No 098433); SP by the Autonomous Province of Trento (“Grandi Progetti 2012, Characterizing and Improving Brain Mechanisms of Attention–ATTEND”); PGS by the Wellcome Trust (Senior Investigator Grant 107802/Z/15/Z). Competing interests: none.

## Additional information

### Author contributions

Conceptualization: CK; Methodology: BLG, RAAI, JG, SP, PGS, CK; Software: BLG, RAAI, CK; Validation: BLG, CK; Formal Analysis: BLG, CK; Investigation: BLG, CK; Resources: BLG, RAAI, CK; Data Curation: BLG; Writing – Original Draft: BLG, CK; Writing – Review & Editing: BLG, RAAI, JG, SP, PGS, CK; Visualization: BLG, CK; Supervision: CK; Project Administration: CK; Funding Acquisition: CK, JG.

---

**Figure S1.**
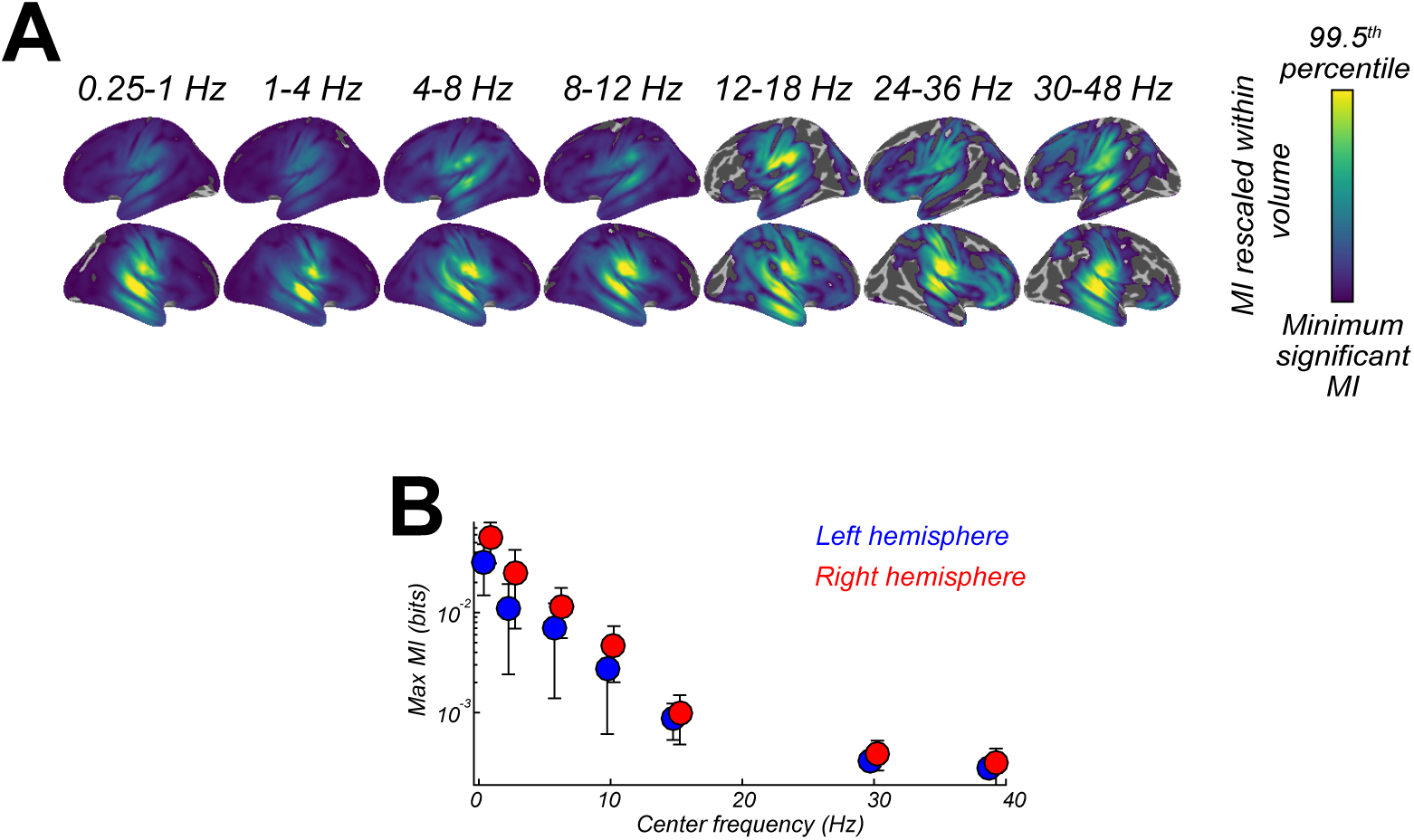
Entrainment of rhythmic MEG activity to the speech envelope. **(A)** Projection of significant speech MI maps, which quantify the entrainment of MEG source activity to the speech envelope, onto the Freesurfer template (FWE = 0.05; proximity = 10 mm; surface-projected significant MI maps rescaled within volume from minimum significant MI to the 99.5^th^ percentile of the surface projection). **(B)** Peak MI in the two hemispheres as a function of frequency (mean ± SEM).

**Figure S2.**
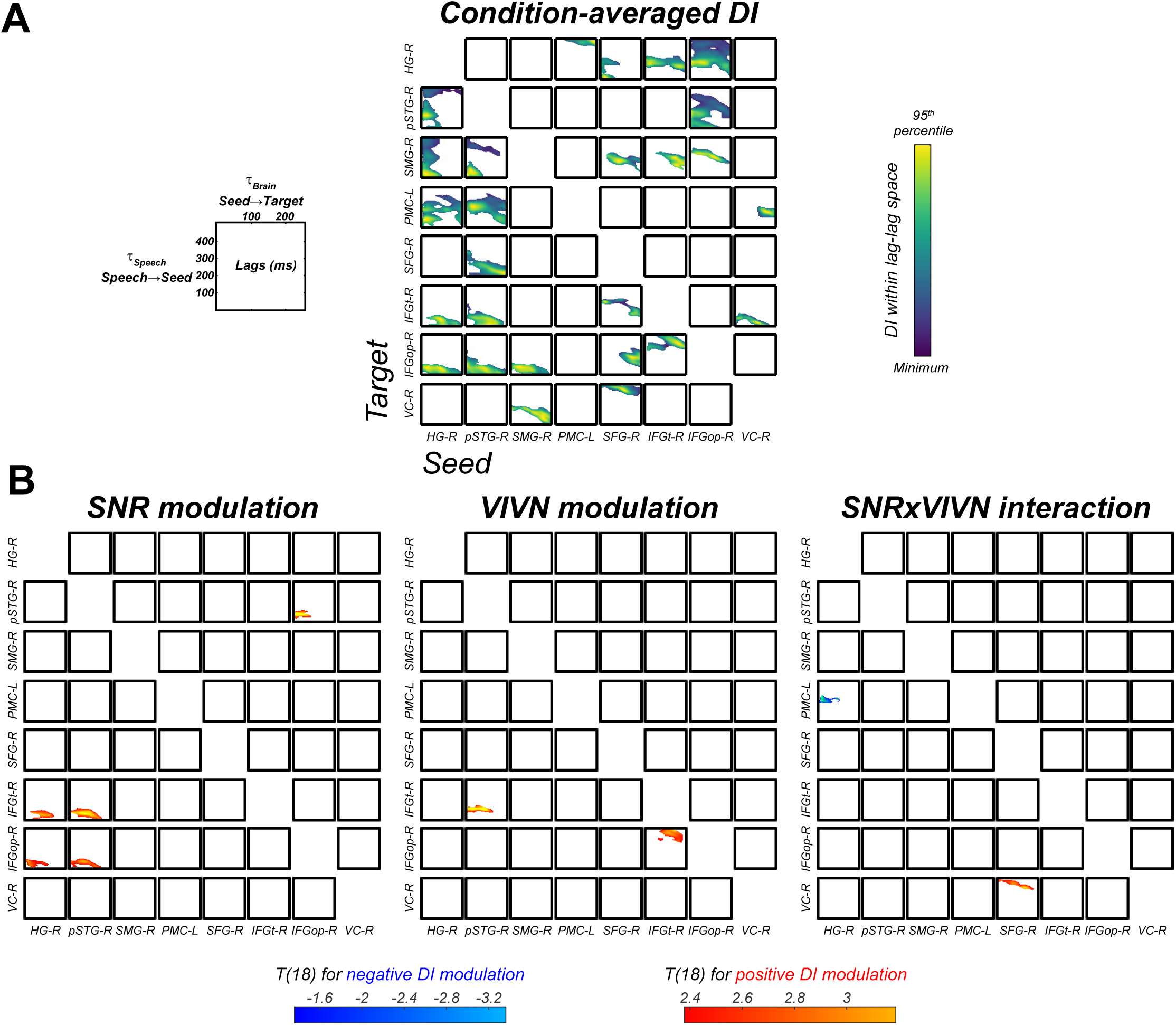
Directed functional connectivity within the speech-entrained network. **(A)** Significant condition-averaged directed information (DI) values between all seed-target pairs as a function of the speech (*τ_Speech_*) and brain lags (*τ_Brain_*). **(B)**: Group-level statistical maps for the GLM effects on DI of acoustic signal quality (SNR), visual informativeness (VIVN) and their interaction.

## References

Abrams DA, Ryali S, Chen T, Balaban E, Levitin DJ, Menon V. Multivariate activation and connectivity patterns discriminate speech intelligibility in Wernicke’s, Broca’s, and Geschwind’s areas. Cereb Cortex. 2013; 23:1703–1714.

Alho J, Lin FH, Sato M, Tiitinen H, Sams M, Jääskeläinen IP. Enhanced neural synchrony between left auditory and premotor cortex is associated with successful phonetic categorization. Front Psychol. 2014; 5:394.

Arnal LH, Giraud AL. Cortical oscillations and sensory predictions. Trends Cogn Sci. 2012; 16:390–398.

Arnal LH, Morillon B, Kell CA, Giraud AL. Dual neural routing of visual facilitation in speech processing. J Neurosci. 2009; 29:13445–13453.

Ashburner J, Friston KJ. Unified segmentation. Neuroimage. 2005; 26:839–851.

Bastos AM, Usrey WM, Adams RA, Mangun GR, Fries P, Friston KJ. Canonical microcircuits for predictive coding. Neuron. 2012; 76:695–711.

Beauchamp MS, Argall BD, Bodurka J, Duyn JH, Martin A. Unraveling multisensory integration: Patchy organization within human STS multisensory cortex. Nat Neurosci. 2004; 7:1190–1192.

Bernstein LE, Auer ET, Takayanagi S. Auditory speech detection in noise enhanced by lipreading. Speech Commun. 2004; 44:5–18.

Besle J, Fischer C, Bidet-Caulet A, Lecaignard F, Bertrand O, Giard MH. Visual activation and audiovisual interactions in the auditory cortex during speech perception: Intracranial recordings in humans. J Neurosci. 2008; 28:14301–14310.

Besserve M, Lowe SC, Logothetis NK, Schölkopf B, Panzeri S. Shifts of gamma phase across primary visual cortical sites reflect dynamic stimulus-modulated information transfer. PLoS Biol. 2015; 13:e1002257.

Besserve M, Schölkopf B, Logothetis NK, Panzeri S. Causal relationships between frequency bands of extracellular signals in visual cortex revealed by an information theoretic analysis. J Comput Neurosci. 2010; 29:547–566.

Binder JR, Desai RH, Graves WW, Conant LL. Where is the semantic system? A critical review and meta-analysis of120 functional neuroimaging studies. Cereb Cortex. 2009; 19:2767–2796.

Bornkessel-Schlesewsky I, Schlesewsky M, Small SL, Rauschecker JP. Neurobiological roots of language in primate audition: Common computational properties. Trends Cogn Sci. 2015; 19:142–150.

Bourguignon M, De Tiége X, Op de Beeck M, Ligot N, Paquier P, Van Bogaert P, et al. The pace of prosodic phrasing couples the listener’s cortex to the reader’s voice. Hum Brain Mapp. 2013; 34:314–326.

Brainard DH. The psychophysics toolbox. Spat Vis. 1997; 10:433–436.

Callan DE, Jones JA, Callan A. Multisensory and modality specific processing of visual speech in different regions of the premotor cortex. Front Psychol. 2014; 5:389.

Callan DE, Jones JA, Callan AM, Akahane-Yamada R. Phonetic perceptual identification by native-and second-language speakers differentially activates brain regions involved with acoustic phonetic processing and those involved with articulatory-auditory/orosensory internal models. Neuroimage. 2004; 22:1182–1194.

Callan DE, Jones JA, Munhall K, Callan AM, Kroos C, Vatikiotis-Bateson E. Neural processes underlying perceptual enhancement by visual speech gestures. Neuroreport. 2003; 14:2213–2218.

Chandrasekaran C, Lemus L, Ghazanfar AA. Dynamic faces speed up the onset of auditory cortical spiking responses during vocal detection. Proc Natl Acad Sci USA. 2013; 110:E4668–E4677.

Chandrasekaran C, Trubanova A, Stillittano S, Caplier A, Ghazanfar AA. The natural statistics of audiovisual speech. PLoS Comput Biol. 2009; 5:e1000436.

Chu YH, Lin FH, Chou YJ, Tsai KWK, Kuo WJ, Jääskeläinen IP. Effective cerebral connectivity during silent speech reading revealed by functional magnetic resonance imaging. PLoS One. 2013; 8:e80265.

Clos M, Langner R, Meyer M, Oechslin MS, Zilles K, Eickhoff SB. Effects of prior information on decoding degraded speech: An fMRI study. Hum Brain Mapp. 2014; 35:61–74.

Cogan GB, Poeppel D. A mutual information analysis of neural coding of speech by low-frequency MEG phase information. J Neurophysiol. 2011; 106:554–563.

Ding N, Simon JZ. Neural coding of continuous speech in auditory cortex during monaural and dichotic listening. J Neurophysiol. 2012; 107:78–89.

Ding N, Simon JZ. Adaptive temporal encoding leads to a background-insensitive cortical representation of speech. J Neurosci. 2013; 33:5728–5735.

Du Y, Buchsbaum BR, Grady CL, Alain C. Noise differentially impacts phoneme representations in the auditory and speech motor systems. Proc Natl Acad Sci USA. 2014; 111:7126–7131.

Evans S, Davis MH. Hierarchical organization of auditory and motor representations in speech perception: Evidence from searchlight similarity analysis. Cereb Cortex. 2015; 25:4772–4788.

Fernández LM, Visser M, Ventura-Campos N, Ávila C, Soto-Faraco S. Top-down attention regulates the neural expression of audiovisual integration. Neuroimage. 2015; 119:272–285.

Ferstl EC, Neumann J, Bogler C, Von Cramon DY. The extended language network: A meta-analysis of neuroimaging studies on text comprehension. Hum Brain Mapp. 2008; 29:581–593.

Fonteneau E, Bozic M, Marslen-Wilson WD. Brain network connectivity during language comprehension: Interacting linguistic and perceptual subsystems. Cereb Cortex. 2015; 25:3962–3976.

Freedman D, Lane D. A nonstochastic interpretation of reported significance levels. J Bus Econ Stat. 1983; 1:292–298.

Ghazanfar AA, Maier JX, Hoffman KL, Logothetis NK. Multisensory integration of dynamic faces and voices in rhesus monkey auditory cortex. J Neurosci. 2005; 25:5004–5012.

Ghazanfar AA, Schroeder CE. Is neocortex essentially multisensory? Trends Cogn Sci. 2006; 10:278–285.

Giraud AL, Poeppel D. Cortical oscillations and speech processing: Emerging computational principles and operations. Nat Neurosci. 2012; 15:511–517.

Gow DW, Olson BB. Sentential influences on acoustic-phonetic processing: A Granger causality analysis of multimodal imaging data. Lang Cogn Neurosci. 2016; 31:841–855.

Grant KW, Seitz PF. Measures of auditory-visual integration in nonsense syllables and sentences. J Acoust Soc Am. 1998; 104:2438–2450.

Greenberg S, Carvey H, Hitchcock L, Chang S. Temporal properties of spontaneous speech—a syllable-centric perspective. Journal of Phonetics. 2003; 31:465–485.

Gross J, Hoogenboom N, Thut G, Schyns P, Panzeri S, Belin P, et al. Speech rhythms and multiplexed oscillatory sensory coding in the human brain. PLoS Biol. 2013; 11:e1001752.

Guediche S, Blumstein S, Fiez J, Holt LL. Speech perception under adverse conditions: Insights from behavioral, computational, and neuroscience research. Front Syst Neurosci. 2013; 7:126.

Hasson U, Ghazanfar AA, Galantucci B, Garrod S, Keysers C. Brain-to-brain coupling: A mechanism for creating and sharing a social world. Trends Cogn Sci. 2012; 16:114–121.

Hasson U, Nusbaum HC, Small SL. Brain networks subserving the extraction of sentence information and its encoding to memory. Cereb Cortex. 2007; 17:2899–2913.

Hasson U, Skipper JI, Nusbaum HC, Small SL. Abstract coding of audiovisual speech: Beyond sensory representation. Neuron. 2007; 56:1116–1126.

Heim S, Eickhoff SB, Amunts K. Specialisation in Broca’s region for semantic, phonological, and syntactic fluency? Neuroimage. 2008; 40:1362–1368.

Hickok G. Computational neuroanatomy of speech production. Nat Rev Neurosci. 2012; 13:135–145.

Hickok G, Poeppel D. The cortical organization of speech processing. Nat Rev Neurosci. 2007; 8:393–402.

Hipp JF, Siegel M. Dissociating neuronal gamma-band activity from cranial and ocular muscle activity in EEG. Front Hum Neurosci. 2013; 7:338.

Ince RAA, Giordano BL, Kayser C, Rousselet GA, Gross J, Schyns PG. A statistical framework for neuroimaging data analysis based on mutual information estimated via a Gaussian copula. Hum Brain Mapp. 2016;. http://doi.org/10.1002/hbm.23471.

Ince RAA, Jaworska K, Gross J, Panzeri S, van Rijsbergen NJ, Rousselet GA, et al. The Deceptively Simple N170 Reflects Network Information Processing Mechanisms Involving Visual Feature Coding and Transfer Across Hemispheres. Cereb Cortex. 2016;. http://doi.org/10.1093/cercor/bhw196.

Ince RAA, van Rijsbergen NJ, Thut G, Rousselet GA, Gross J, Panzeri S, et al. Tracing the Flow of Perceptual Features in an Algorithmic Brain Network. Sci Rep. 2015; 5:17681.

Kayser C, Logothetis NK, Panzeri S. Visual enhancement of the information representation in auditory cortex. Curr Biol. 2010; 20:19–24.

Kayser C, Wilson C,Safaai H, Sakata S, Panzeri S. Rhythmic Auditory Cortex Activity at Multiple Timescales Shapes Stimulus-Response Gain and Background Firing. J Neurosci. 2015; 35:7750–7762.

Kayser SJ, Ince RAA, Gross J, Kayser C. Irregular speech rate dissociates auditory cortical entrainment, evoked responses, and frontal alpha. J Neurosci. 2015; 35:14691–14701.

Kayser SJ, McNair SW, Kayser C. Prestimulus influences on auditory perception from sensory representations and decision processes. Proc Natl Acad Sci USA. 2016; 113:4842–4847.

Keitel A, Ince RAA, Gross J, Kayser C. Auditory cortical delta-entrainment interacts with oscillatory power in multiple fronto-parietal networks. Neuroimage. 2017; 147:32–42.

Krieger-Redwood K, Gaskell MG, Lindsay S, Jefferies E. The selective role of premotor cortex in speech perception: A contribution to phoneme judgements but not speech comprehension. J Cognitive Neurosci. 2013; 25:2179–2188.

Kriegeskorte N, Goebel R, Bandettini P. Information-based functional brain mapping. Proc Natl Acad Sci USA. 2006; 103:3863–3868.

Lakatos P, O’Connell MN, Barczak A, Mills A, Javitt DC, Schroeder CE. The leading sense: Supramodal control of neurophysiological context by attention. Neuron. 2009; 64:419–430.

Lee H, Noppeney U. Physical and perceptual factors shape the neural mechanisms that integrate audiovisual signals in speech comprehension. J Neurosci. 2011; 31:11338–11350.

Maris E, Oostenveld R. Nonparametric statistical testing of EEG and MEG data. J Neurosci Methods. 2007; 164:177–190.

Massey J. Causality, feedback and directed information. In: Proc Int Symp Inf Theory Applic (ISITA-90); 1990. p. 303–305.

McGettigan C, Faulkner A, Altarelli I, Obleser J, Baverstock H, Scott SK. Speech comprehension aided by multiple modalities: Behavioural and neural interactions. Neuropsychologia. 2012; 50:762–776.

Meister IG, Wilson SM, Deblieck C, Wu AD, Iacoboni M. The essential role of premotor cortex in speech perception. Curr Biol. 2007; 17:1692–1696.

Mesgarani N, Chang EF. Selective cortical representation of attended speaker in multi-talker speech perception. Nature. 2012; 485:233–236.

Morillon B, Hackett TA, Kajikawa Y, Schroeder CE. Predictive motor control of sensory dynamics in auditory active sensing. CurrOpin Neurobiol. 2015; 31:230–238.

Nath AR, Beauchamp MS. Dynamic changes in superior temporal sulcus connectivity during perception of noisy audiovisual speech. J Neurosci. 2011; 31:1704–1714.

Ng BSW, Logothetis NK, Kayser C. EEG phase patterns reflect the selectivity of neural firing. Cereb Cortex. 2013; 23: 389–398.

Ng BSW, Schroeder T, Kayser C. A precluding but not ensuring role of entrained low-frequency oscillations for auditory perception. J Neurosci. 2012; 32:12268–12276.

Oostenveld R, Fries P, Maris E, Schoffelen JM. FieldTrip: Open source software for advanced analysis of MEG, EEG, and invasive electrophysiological data. Comput Intell Neurosci. 2010; 2011:156869.

Osnes B, Hugdahl K, Specht K. Effective connectivity analysis demonstrates involvement of premotor cortex during speech perception. Neuroimage. 2011; 54:2437–2445.

Panzeri S, Senatore R, Montemurro MA, Petersen RS. Correcting for the sampling bias problem in spike train information measures. J Neurophysiol. 2007; 98:1064–1072.

Park H, Ince RAA, Schyns PG, Thut G, Gross J. Frontal top-down signals increase coupling of auditory low-frequency oscillations to continuous speech in human listeners. Curr Biol. 2015; 25:1649–1653.

Park H, Kayser C, Thut G, Gross J. Lip movements entrain the observers’ low-frequency brain oscillations to facilitate speech intelligibility. eLife. 2016; 5:e14521.

Peelle JE, Davis MH. Neural oscillations carry speech rhythm through to comprehension. Front Psychol. 2012; 3:320.

Peelle JE, Sommers MS. Prediction and constraint in audiovisual speech perception. Cortex. 2015; 68:169–181.

Pickering MJ, Garrod S. An integrated theory of language production and comprehension. Behav Brain Sci. 2013; 36:329–347.

Poeppel D. The analysis of speech in different temporal integration windows: Cerebral lateralization as “asymmetric sampling in time”. Speech Commun. 2003; 41:245–255.

Poeppel D. The neuroanatomic and neurophysiological infrastructure for speech and language. CurrOpin Neu-robiol. 2014; 28:142–149.

Price CJ. A review and synthesis of the first 20 years of PET and fMRI studies of heard speech, spoken language and reading. Neuroimage. 2012; 62:816–847.

Rauschecker JP, Scott SK. Maps and streams in the auditory cortex: Nonhuman primates illuminate human speech processing. Nat Neurosci. 2009; 12:718–724.

Riedel P, Ragert P, Schelinski S, Kiebel SJ, von Kriegstein K. Visual face-movement sensitive cortex is relevant for auditory-only speech recognition. Cortex. 2015; 68:86–99.

Rohe T, Noppeney U. Cortical hierarchies perform Bayesian causal inference in multisensory perception. PLoS Biol. 2015; 13:e1002073.

Ross LA, Saint-Amour D, Leavitt VM, Javitt DC, Foxe JJ. Do you see what I am saying? Exploring visual enhancement of speech comprehension in noisy environments. Cereb Cortex. 2007; 17:1147–1153.

Schepers IM, Yoshor D, Beauchamp MS. Electrocorticography reveals enhanced visual cortex responses to visual speech. Cereb Cortex. 2015; 25:4103–4110.

Schreiber T. Measuring information transfer. Phys Rev Lett. 2000; 85:461–464.

Schroeder CE, Lakatos P, Kajikawa Y, Partan S, Puce A. Neuronal oscillations and visual amplification of speech. Trends Cogn Sci. 2008; 12:106–113.

Schwartz JL, Berthommier F, Savariaux C. Seeing to hear better: Evidence for early audio-visual interactions in speech identification. Cognition. 2004; 93:B69–B78.

Schwartz JL, Savariaux C. No, there is no 150 ms lead of visual speech on auditory speech, but a range of audiovisual asynchronies varying from small audio lead to large audio lag. PLoS Comput Biol. 2014; 10:e1003743.

Skipper JI, Goldin-Meadow S, Nusbaum HC, Small SL. Gestures orchestrate brain networks for language understanding. Curr Biol. 2009; 19:661–667.

Sumby WH, Pollack I. Visual contribution to speech intelligibility in noise. J Acoust Soc Am. 1954; 26:212–215.

van Atteveldt N, Formisano E, Goebel R, Blomert L. Integration of letters and speech sounds in the human brain. Neuron. 2004; 43:271–282.

van Wassenhove V. Speech through ears and eyes: Interfacing the senses with the supramodal brain. Front Psychol. 2013; 4:2.

VanderGhinst M, Bourguignon M, Op de Beeck M, Wens V, Marty B, Hassid S, et al. Left Superior Temporal Gyrus Is Coupled to Attended Speech in a Cocktail-Party Auditory Scene. J Neurosci. 2016; 36:1596–1606.

Vetter P, Smith FW, Muckli L. Decoding sound and imagery content in early visual cortex. Curr Biol. 2014; 24: 1256–1262.

Vicente R, Wibral M, Lindner M, Pipa G. Transfer entropy-a model-free measure of effective connectivity for the neurosciences. J Comput Neurosci. 2011; 30:45–67.

Wibral M, Rahm B, Rieder M, Lindner M, Vicente R, Kaiser J. Transfer entropy in magnetoencephalographic data: Quantifying information flow in cortical and cerebellar networks. Prog Biophys Mol Biol. 2011; 105:80–97.

Wild CJ, Yusuf A, Wilson DE, Peelle JE, Davis MH, Johnsrude IS. Effortful listening: The processing of degraded speech depends critically on attention. J Neurosci. 2012; 32:14010–14021.

Wilson SM, Saygin AP, Sereno MI, Iacoboni M. Listening to speech activates motor areas involved in speech production. Nat Neurosci. 2004; 7:701–702.

Winkler AM, Ridgway GR, Webster MA, Smith SM, Nichols TE. Permutation inference for the general linear model. Neuroimage. 2014; 92:381–397.

Wright TM, Pelphrey KA, Allison T, McKeown MJ, McCarthy G. Polysensory interactions along lateral temporal regions evoked by audiovisual speech. Cereb Cortex. 2003; 13:1034–1043.

Yarkoni T, Speer NK, Zacks JM. Neural substrates of narrative comprehension and memory. Neuroimage. 2008; 41:1408–1425.

Zion Golumbic E,Cogan BG, Schroeder CE, Poeppel D. Visual input enhances selective speech envelope tracking in auditory cortex at a “cocktail party”. J Neurosci. 2013; 33:1417–1426.

